# Probiotic strains improve high-fat diet-induced hypercholesterolemia through modulating gut microbiota in ways different from atorvastatin

**DOI:** 10.1101/712083

**Authors:** Sudun, Songling Liu, Chen Xiao, Can Peng, Lifeng Liang, Xiaofen He, Shancen Zhao, Gengyun Zhang

## Abstract

Hypercholesterolemia is a major risk factor for cardiovascular disease (CVD). Probiotics is one of the most popular dietary supplements for hypercholesterolemia, but there are questions as to whether there are differences between probiotics and cholesterol-lowering drugs as like atorvastatin (ATO) both in effectiveness and the underlying mechanisms. In this study, the hypocholesterolemia effects of 4 probiotic strains were investigated and compared with ATO, focusing on their impacts on gut microbiota. Hypercholesterolemia model was established via high-fat diet (HFD) in golden hamsters after which ATO and the 4 probiotics were orally administered individually for 8 weeks. All probiotics were effective, but less than ATO, both on body weight, serum parameters (TG, TC, LDL, INS, HbA1c) and expression of inflammatory factors (TNF-α, IL-1β, CRP), with strain JQII-5 most significant. Besides, these effects were associated with restoration of the microbiota dysbiosis induced by HFD. It was worth noting that ATO and probiotics induced different shifts of gut microbiota in both structure and key phylotypes. Most interestingly, Allobaculum, a HFD-suppressed genus, reported to be involved in alleviating oxidative stress, was enriched by all tested probiotic strains, but not by ATO. Furthermore, Prevotella, also a HFD-suppressed genus, was uniquely reversed by JQII-5. Importantly, most of the alerted genus and reversed genus was found to be correlated to inflammatory state and serum lipid level. Compared with ATO, probiotic strains were less effective on body weight, hypercholesterolemia, and inflammation. However, probiotics exert additional favorable effects on gut microbiota, making them excellent potential complements to cholesterol-lowering drugs like ATO.

## 1. Introduction

Hypercholesterolemia is a major risk factor for the development of cardiovascular disease (CVD).^1^ It was reported that hypercholesterolemia contributed to 45% of heart attacks in Western Europe from 1999-2003.^2^ Therefore, it is not surprising that hypercholesterolemia has received particular attention from both societies and scientists. To effectively control the development of hypercholesterolemia, both drugs and dietary supplements were exploited. However, many cholesterol-lowering drugs present important side effects, restricting their long-term administration. For this reason, dietary control or supplementation were preferred as complementary therapies, of which probiotics are the one of the most notable.^3^

At present, although many probiotic strains have been demonstrated to exert significantly beneficial effects on lowering LDL and total cholesterol (TC),^4-6^ there is still not enough evidence to confirm their clinical efficiency. For one reason, controversial results have been reported attributing to various external factors, such as strains differences, administration dosage, clinical characteristic of subjects, and duration of treatment period.^7-9^ Moreover, studies examining the efficacy of probiotics in reducing cholesterol often do not sufficiently address the mechanism by which they take effects, and which is extensively explicit when a cholesterol-lowering drug was exploited.^3^A comprehensive comparation of probiotic strains with a well-proved cholesterol-lowering drug would provide new insights into the effective use of probiotic strains in the improvement of hypercholesterolemia.

High-fat diet (HFD) induced alteration of gut microbiota has been demonstrated to play a crucial role in the development of hypercholesterolemia. ^10, 11^ And with the development of research, increasing evidence has demonstrated that the gut microbiome was associated not only with the pathogenesis of disease, but also with the effects of disease treatment and prevention. ^12, 13^ Accordingly, targeting the gut microbiota using dietary interventions, prebiotics and probiotics have been shown to prevent or alleviate the metabolic diseases in the past few years. ^14^ Furthermore, many drugs were investigated for their effects on gut microbiota in recent years. ^15, 16^ Among these, atorvastatin (ATO) is a well proved and commonly prescribed cholesterol-lowering drug. It is an inhibitor of 3-hydroxy-3-methyglutaryl-coenzyme A (HMG-CoA) reductase. This enzyme catalyzes the conversion of HMG-CoA to mevalonate, an early and rate-limiting step in cholesterol biosynthesis.^17^ Especially, ATO has been recently reported to reverse the alteration of some specific dominant taxa induced by HFD in rats.^15^

In this study we compared the hypocholesterolemia effects of 4 cholesterol-degrading probiotics: *Pediococcus acidilactici* JQII-5 (JQII-5), *Pediococcus pentosaceus* JQI-7(JQI-7), *Lactobacillus plantarum* LLY-606 (LLY-606) and *Lactobacillus plantarum* PC-26,^18^ with ATO, and investigated the mechanisms by which they exerted hypochelesterolemic effects, focusing on their impact on gut microbiota.

## 2. Materials and Methods

### 2.1 Bacterial strains

*Pediococcus acidilactici* JQII-5 (JQII-5) and *Pediococcus pentosaceus* JQI-7(JQI-7) were isolated from Traditional yoghurt from Inner Mongolia; *Lactobacillus plantarum* LLY-606 (LLY-606) and *Lactobacillus plantarum* PC-26 was isolated from Human Intestinal Tract. They were all stored at China General Microbiological Culture Collection Center with a preservation number of CGMCC No. 10512 for JQII-5, CGMCC No. 10511 for JQI-7, CGMCC No. 13984 for LLY-606 and CGMCC No. 12810 for pc-26 respectively. The strains of JQII-5, JQI-7, LLY-606 and pc-26 are respectively cultivated in Man-Rogosa-Sharpe (MRS) broth at 37°C for 24h. After the incubation, the LABs pellet was collected by centrifugation at 12000 g and 4°C for 15 min, and washed twice with cold sterile water. The LABs pellet was finally lyophilized and stored at-80°C for further experiment. The lyophilized LABs were dissolved in distilled water to a concentration of 10^8^ CFU/mL before use.

### 2.2 Animals and Experiment Design

All interventions and animal care procedures were carried in accordance with the Guidelines and Policies for Animal Surgery of the Institute of Medical Plant Development, Chinese Academy of Medical Sciences & Peking Union Medical College (Beijing, China) and approved by the institutional Animal Use and Care Committee. Seventy specific pathogen-free male golden hamsters aged 6-8 weeks old and weighing 120 ± 20 g were purchased from the Vital River Laboratory Technology Co., Ltd. (Beijing, China), and were housed in stainless steel cages in a temperature-controlled room (22-25°C, 50% to 70% relative humidity) with a 12 h light/dark cycle (light on 7:00 AM), and distilled deionized water was provided continuously. After acclimatization for 1 week, ten hamsters were selected randomly as the normal control diet group (NCD, n = 10) and fed with a normal diet throughout the whole experimental period. Others were fed with a high-fat diet for 2 weeks to generate hypercholesterolemic model. The high-fat diet contained 15% (w/w) lard oil and 0.2% cholesterol and 84.8% normal diet (Beijing HuaFuKang Biotechnology Co., Ltd., Beijing, China).

After the hypercholesterolemic model was successfully established (0th week of probiotic intervention), sixty hyperlipidemic hamsters were randomly divided into the following six groups (n = 10/each group): (1) HFD: hamsters on high-fat diet; (2) ATO: hamsters on high-fat diet plus atorvastatin (Pfizer Ltd., Dalian, China); (3) JQII-5: hamsters on high-fat diet plus *Pediococcus acidilactici* JQII-5; (4) JQI-7: hamsters on high-fat diet plus *Pediococcus pentosaceus* JQI-7; (5) LLY-606: hamsters on high-fat diet plus *Lactobacillus plantarum* LLY-606; (6) pc-26: hamsters on high-fat diet plus *Lactobacillus plantarum* pc-26. The atorvastatin was diluted by water at 1 mg/mL using a magnetic stir and was orally administered to ATO group with a dose of 3mg/kg/day. All probiotic strains were administrated at a dose of 10^8^cfu daily, the NCD and HFD group received the same volume of distilled water by oral administration once a day for eight weeks. The dose of administration of ATO and strains were determined by referring to previous studies and was verified by preliminary experiments. ^6,15^

### 2.3 Preparation of Blood Samples

Blood samples were obtained from the retro-orbital sinus into prechilled tubes on 0th, 4th and 8th week of probiotic intervention. The serum was separated by centrifugation at 3000 g 4°C for 10 min and stored at −80°C for future analyses.

### 2.4 Analysis of Serum Lipids

Concentrations of serum lipids including TC, triglycerides (TG), HDL-C and LDL-C were determined on 0th, 4th and 8th week using assay kits (BioSino Bio-Technology & Science Inc., Beijing, China) and the AU480 Chemistry analyzer (Beckman Coulter Inc., California, USA).

### 2.5 Inflammation Cytokine Quantification in Serum

Serum concentrations of tumor necrosis factor-α (TNF-α) and interleukin1β (IL-1β) were measured on 8th week using Valukine™ immunoassay kits (R&D Systems, Inc., Minneapolis, USA) according to the manufacturer’s instructions using microplate reader (Synergy 2, BioTek, USA). Serum C-reactive protein (CRP) level was determined using radioimmunoassay kit (YingHua Biotechnology Research Institute, Beijing, China) using Radioimmunocounter (XH-6020, Xian Nuclear instrument Ltd., China) on 8th week.

### 2.6 Measurement of Glycometabolism indices

Serum insulin (INS) concentration was measured using radioimmunoassay kits (YingHua Biotechnology Research Institute, Beijing, China) using Radioimmunocounter (XH-6020, Xian Nuclear instrument Ltd., China) on 8th week. The quantitative determination of glycated hemoglobin A1c (HbA1c) was done according to the procedures in a commercial kit (Nanjing Jiancheng Biotechnology Institute, JiangSu, China) using microplate reader (Synergy 2, BioTek, USA).

### 2.7 Gut Microbiota Analysis

Fresh fecal samples were collected from each group on 8th week and stored at −80°C for DNA extraction. Microbial genome DNA from the fecal samples was extracted using QIAamp DNA stool mini kit (Qiagen Inc., Hilden, Germany) according to the manufacture’s recommendation. The V4 hypervariable region of the 16S rDNA was PCR amplified from the microbial genome DNA which were harvested from fecal samples using universal primer (forward primers: 5’-AYTGGGYDTAAAGNG-3’, recerse primers: 5’-TACNVGGGTATCTAATCC-3’). The PCR condition were 94°C for 5 min, followed by 30 cycles of 94°C for 30 s, 58°C for 30 s (annealing) and 72°C for 30 s (extension), and then 72°C for 5 min.

The PCR products were excised from a 1.5% agarose gel and purified by AxyPrep Gel Extraction Kit (Axygen, Scientific Inc., Union City, CA, USA). They were then quantified by PicoGreen dsDNA Assay Kit (Life Technologies Inc., Grand Island., NY, USA) and BioTek Microplate Reader (BioTek Inc., Winooski, VT, USA). Barcoded V4 amplicon was sequenced using the paired-end method by Illumina Miseq at BGI TechSolutions Co., Ltd (Wuhan, China). Sequences reads with an average quality score lower than 20, ambiguous bases, homopolymer > 7 bases, containing primer mismatches, or reads length shorter than 100 bp were removed. For V4 paired read, only sequences that overlapped longer than 10 bp and without any mismatches were assembled (Tian, Ma et al. 2013). Reads which could not be assembled were removed. Sequences analysis were performed by Uparse (v7.0.1090) and sequences with ≥ 97% similarity were assigned to the same operational taxonomic units (OTUs). For each representative sequence for each OUT, the GreenGene Database was used based on RDP classifer (v2.2) to annotate taxonomic information. The differences of the dominant species in different groups and multiple sequence alignment were conducted using the MUSLE software to study phylogenetic relationship of different OUTs. Alpha diversity and beta diversity were calculated by QIIME software (v1.80) to evaluate species diversity. Sequence data of this study is available in China Nucleotide Sequence Archive under project No. CNP0000290.

### 2.8 Determination of SCFAs in Feces

SCFAs were measured according to the method of Wang *et al* with modifications. ^19^ 0.3g of stool sample were homogenized with a Mini Bead-heater-16 (BioSpec Products, Inc, Bartlesville, OK, USA) in 1mL distilled water. Samples were centrifuged at 12000g for 10min. 800ul supernatant was collected. Then precipitation was rehomogenized in 800uL distilled water and was centrifuged at 12000g for 10min. 700ul supernatant was collected at this time. All together 1500ul supernatant were collected, and these were pooled for SCFA analysis.

SCFA levels were measured by high performance liquid chromatograph (LC-2030, Shimadzu Corporation, Japan). Supernatants (20uL) were separated isocrattically at a flow rate of 1mL/min at 65 °C on a 250-by 4.6-mm YMC-Pack Pro C18 column (HanKing Instrument & Equipment, Shanghai, China), using 0.025% phosphoric acid and 5% acetonitrile as the mobile phase. The identification and quantification were performed at 210nm. Standard solutions of lactic, acetic and propionate were run separately for identification and quantification of each compound.

### 2.9 Statistical Analysis

The biodiversity and richness of OUTs were calculated using QIME 1.9 implemented with nonparametric Chao1 and rarefaction analysis. Data was normalized to an equal number of reads per sample, and PCA was performed from the sequences at OUT level with >97% similarity using unsupervised multivariate statistical method. Nonparametric factorial Kruskal-Wallis rank sum test was used were performed to identify significantly different bacterial taxa among different groups. Differences in body weight, cholesterol levels, inflammatory levels and short fatty acid levels were analyzed statistically using IBM SPSS Statistics 22.0 software. The data values are presented as the Mean ± SD. One-way analysis of variance (ANOVA) with Duncan’s test was used to measure the significance of the main effects (*p* < 0.05) in the variable between groups. Correlations level were calculated using IBM SPSS Statistics 22.0 software.

## 3. Results

### 3.1 Effects of probiotics and ATO on gain of body weight induced by HFD

Prior to administration ATO or probiotics, 2 weeks of HFD feeding had induced significant increase in the body weight compared with normal chow diet (NCD) group (Figure 1). After 3 weeks co-administration of ATO, the increase of body weight observed in HFD-ATO group had been prevented. In fact, animals in this group displayed no significant difference in body weight from those in NCD group. However, it was not until the 4th week, the increase in body weight of groups HFD+JQII-5, HFD+JIQ-7, HFD+LLY-606, and HFD+PC-26 started to be attenuated. Even so, the body weight of animals in all these probiotic groups was still significantly higher than that in both HFD+ATO and NCD groups until the 8th week.

**Figure 1.**
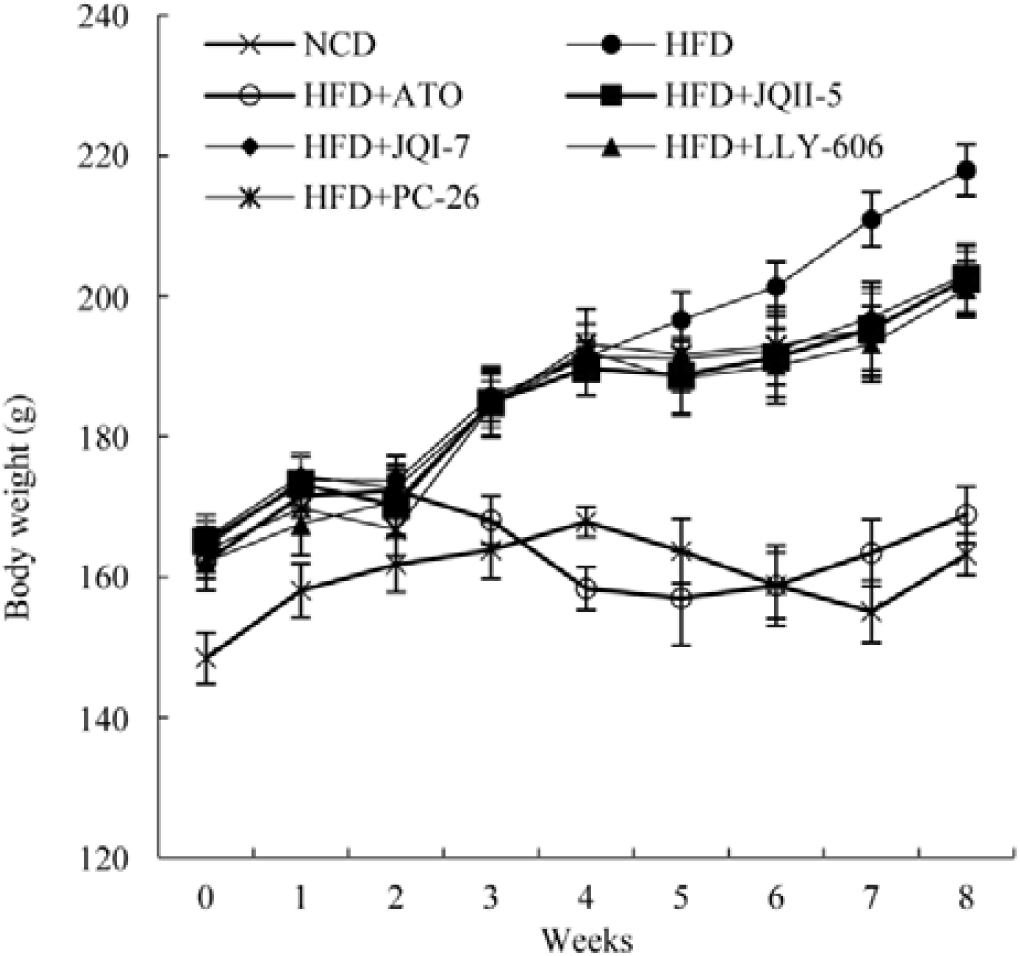
Effects of ATO and probiotic strains on the body weight of golden hamsters.

### 3.2 Hypocholesterolemic effects of ATO and probiotic strains

The concentration of serum TC, TG, HDL-C and LDL-C in all groups were assayed at 0th, 4th and 8th week of co-administration. HFD feeding induced significant higher TG, TC and LDL-C compared with NCD (*p* < 0.05), which indicated that 4 weeks of co-administration of ATO significantly decreased TC, TG and LDL level compared with HFD, and these effects were maintained until the 8th week. However, all the probiotic co-administration groups did not significantly decreased TC or LDL-C level after 4 weeks intervention (**Figure 2B and Figure 2C**). Nonetheless, 4 weeks of co-administration of probiotic prevented the increase of TG level observed in HFD groups, although it was still significantly higher than in HFD+ATO and NCD groups (**Figure 2A**).

**Figure 2.**
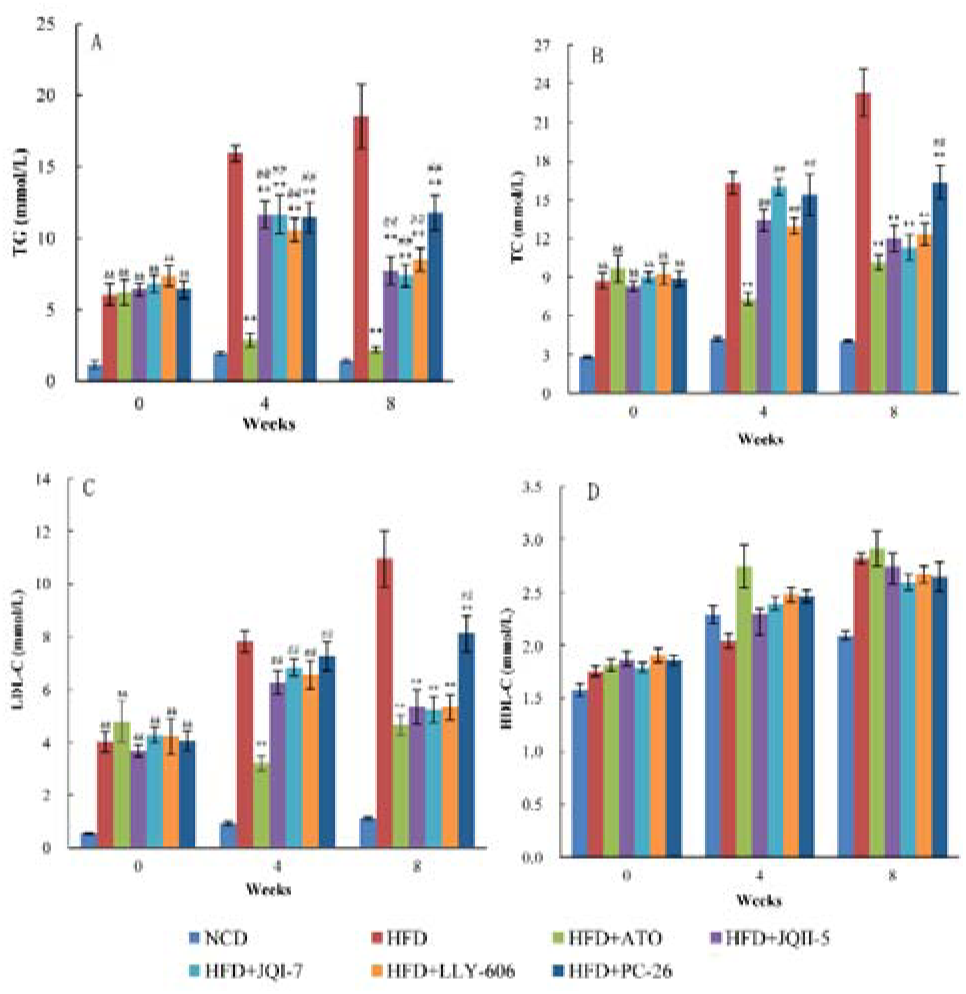
Effects of ATO and probiotics on high-fat diet induced hypercholesterolemia of golden hamsters. A: Serum TG levels; B: Serum TC levels; C: Serum HDL-C levels; D: Serum LDL-C levels. ^&&^means significant difference fron NCD group. ** means significant difference from HFD group; ^##^means different differnce from ATO Group (*p* < 0.05, n = 10).

At the 8th week, the TC level of all probiotic groups started to be significantly lower than the HFD group and to show no difference from the HFD+ATO group. The TG level of all probiotic groups was also decreased to be significantly lower than HFD, but it was still higher than the HFD+ATO (**Figure 2A**). The effects on the LDL-C was divergent among the probiotic groups at the 8th week of co-administration. Although co-administration of JQII-5, JIQ-7 or LLY-606 decreased the concentration of LDL-C to level comparable to HFD+ATO, it was still higher in HFD+PC-26 than in HFD+ATO (**Figure 2C**).

These results showed that compared with ATO, probiotic strains used in this study showed mild effectiveness on hypercholesterolemia. Further, the strains of JQII-5, JIQ-7 and LLY-606 were more significant than PC-26 in the improvement of hypercholesterolemia, especially on reducing the concentration of LDL-C.

### 3.3 Effects of probiotics and ATO on expression of inflammatory factors

After 8 weeks of co-administration of ATO or probiotics, the expression of inflammatory cytokines was assayed. Just as expected, high-fat diet feeding was accompanied with increased TNF-α (*p* < 0.05), IL-1β (*p* < 0.05) and CPR (*p* < 0.05) (**Figure 3**). Co-administration of ATO significantly decreased the TNF-α, IL-1β and CRP. In all the four probiotic groups, IL-1β was decreased to the level which was significantly lower than that in HFD group, but higher than in FHD+ATO group (**Figure 3A**). At the same time, co-administration of JQII-5, PC-26 also induced a significantly lower expression of TNF-α (*p* < 0.05) compared with HFD, but co-administration of JIQ-7 and LYY-606 did not decrease TNF-α (**Figure 3B**). The CRP concentration of the HFD+JQII-5 and HFD+JQI-7 groups was significantly lower (*p* < 0.05) than HFD and it was comparable to that in group HFD+ATO (**Figure 3C**).

**Figure 3.**
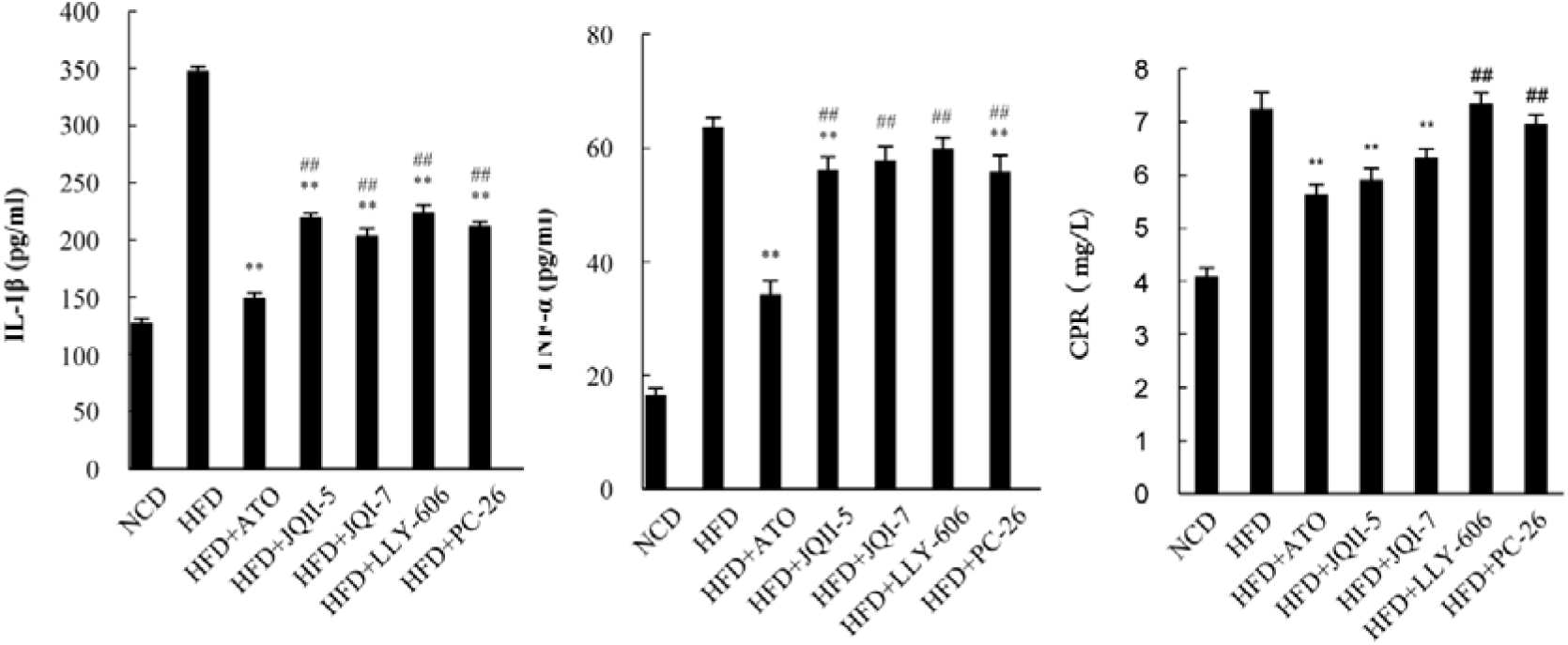
Effects of ATO and probiotics on the expression inflammatory cytokine of golden hamsters. A: IL-1β; B: TNF-α; C: CPR. **means significant difference from HFD group; ^##^means different differnce from ATO Group (*p* < 0.05, n = 10).

The probiotic strains in this study were proved to be able to reduce the expression inflammatory factors induced by high-fat diet, although their effects were not as remarkable as ATO and were divergent among themselves, with JQII-5 and JQI-7 more potential than LLY-606 and PC-26.

### 3.4 Effect on probiotics and ATO on Glycometabolism

Compared with NCD group, feeding high-fat diet significantly increased (*p* < 0.05) INS and HbAc1 concentration, and administration of ATO successfully decreased both INS and HbAc1 (**Figure 4**). Administration of JQII-5 significantly decreased the INS and HbAc1 to levels comparable to ATO (**Figure 4A** and **Figure 4B**). HFD+JIQ-7 group also showed significantly lower concentration of HbAc1 and INS compared with HFD group, but not as low as HFD+ATO group. LLY-606 and pc-26 significantly decreased HbAc1 compared with HFD, but not to the level comparable with ATO.

**Figure 4.**
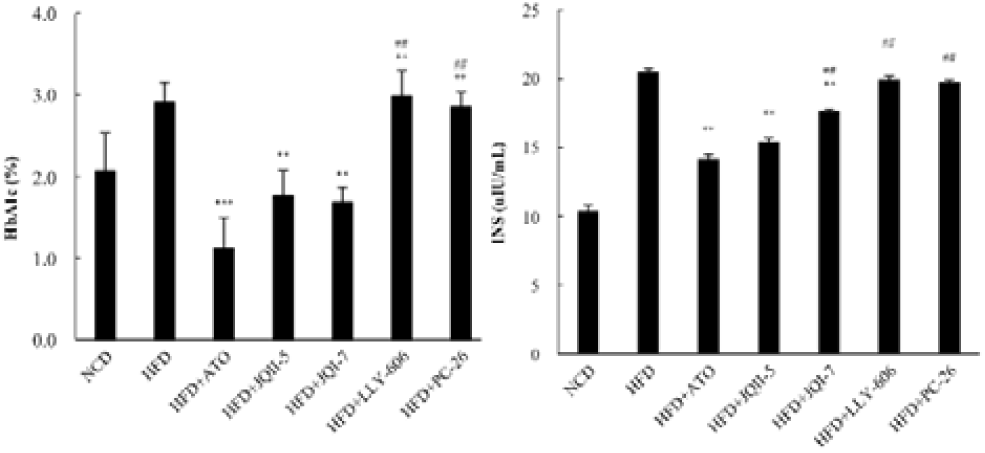
Effects of ATO and probiotics on the glycometabolism of golden hamsters. A :HbA1c; B:INS; **means significant difference from HFD group; ^##^means different differnce from ATO Group (*p* < 0.05, n = 10).

As for the glcometabolism index, the effects of different probiotic strains exhibited more distinct divergence, with JQII-5 more effective than JIQ-7, LLY-606 and PC-26.

### 3.5 Effect of Probiotics and ATO on Gut Microbiota

#### 3.5.1 Richness and Diversity Indices

After spliced and optimized, 70 samples were delineated into 1358 OTUs at the 97% similarity level with distance-based OTU and richness. The richness and diversity indices of gut microbiota in different groups were listed in Table 1. ATO co-administration did not influence the diversity. The observed species and Chao indices of the HFD+PC-26 group were significantly lower (p < 0.05) than the HFD+JQI-7 and HFD+LYY-606 group and showed no difference from the HFD+JQII-5 group. The ACE and Shannon indices showed no difference among the four single-strain groups. At the same time, the Simpson index of the HFD+JQII-5 and HFD+LYY-606 group was significantly lower (p < 0.05) than the HFD+JQI-7 group and showed no difference from the HFD+PC-26 group.

**Table 1.** The OTU richness and diversity indices of different groups.

#### 3.5.2 Overall structural change of gut microbiota in response of ATO or probiotics

The overall structural changes of the gut microbiota were analyzed using unsupervised multivariate statistical method of PCA. PCA scores plot showed that the gut microbiota of the HFD group present a structural shift from the NCD group (*Figure 5*). As the figure 5 showed, the communities of all the probiotic groups present a structural shift from the ATO group, and the communities among JQI-7, LLY-606 and PC-26 showed more similarity than they did with JQII-5 group.

**Figure 5.**
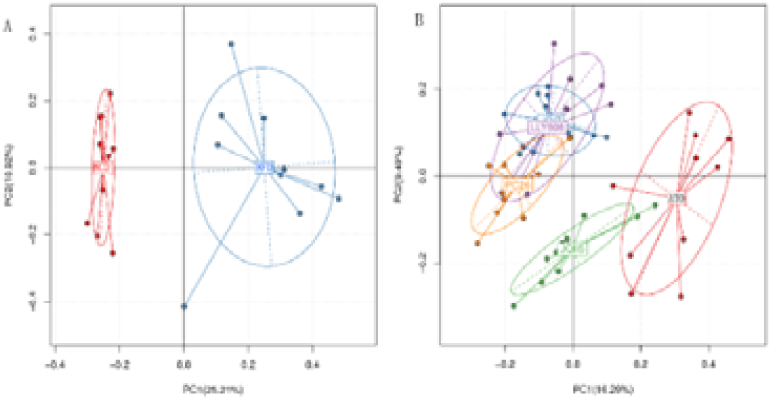
Effects of ATO and probiotics on structure of gut microbiota analyzed by principal composition analysis (PCA). A:Comparation between HFD and NCD; B : comparation among HFD+ATO(ATO),HFD+JQII-5(JQII5),HFD+JQI-7 (JQI7),HFD+LLY-606(LYY606) and HFD+PC-26(PC26)

HFD significantly increased the relative abundance of the Bacteroidetes (*p*<0.01) (*table 2*). What is the not expected is that JQII-5 did not change the structure at the phylum level. Except to JQII-5, all the other probiotic and ATO co-administration reversed the shifts in Bacteroidetes and Firmicutes.

**Table 2.**
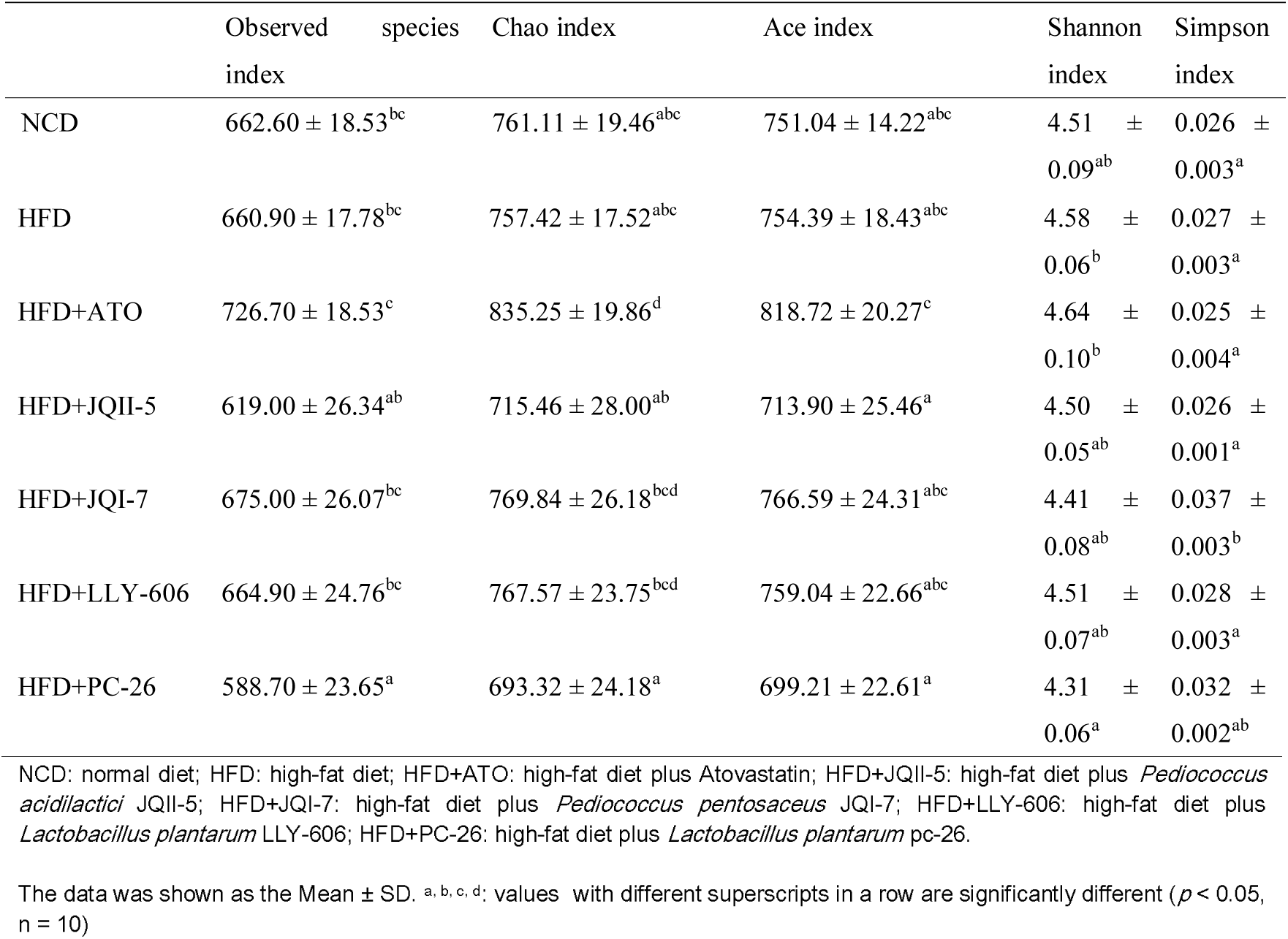
The OTU richness and diversity indices of different groups.

**Table 3.**
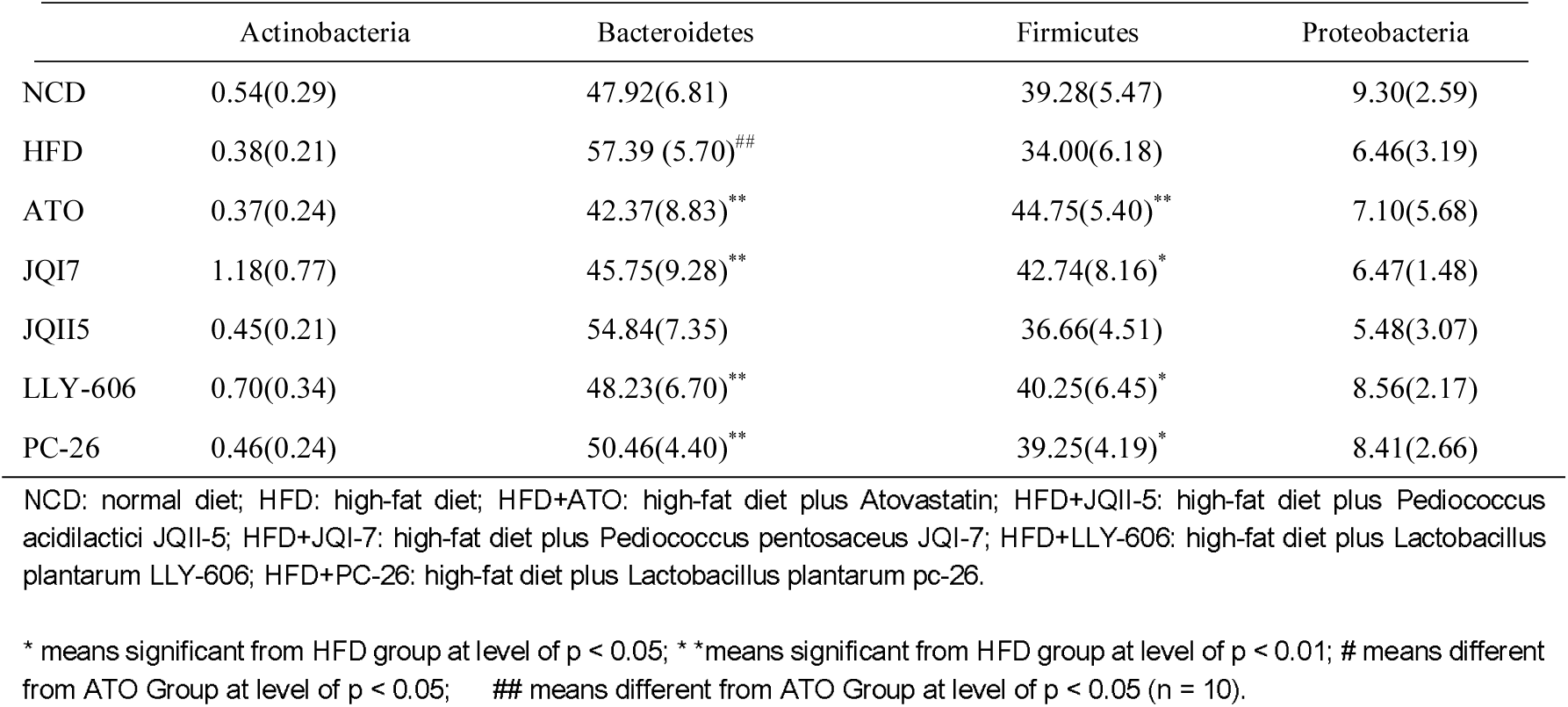
Effects of ATO and probiotics on the structure of gut microbiota at the phylum level

### 3.6 Key phylotypes of gut microbiota modulated by ATO or probiotics

To identify key phylotypes of the gut microbiota responding to HFD, and to compared the key phylotypes response to ATO, JQII-5, JQI-7, LLY-606 and PC-26, nonparametric factorial Kruskal-Wallis rank sum test was used. Linear discriminant analysis scores higher than 2 was considered a higher abundance in the corresponding group than the other group. The relative abundance of significantly different species was showed by linear discriminant analysis scores effect size (LEfSe) taxonomy cladogram (supplementary S1-S5).

First of all, the NCD and FHD groups were compared to find the HFD induced changes in gut microbiota. At the genus level, HFD showed significant selective suppression of Allobaculum, Ruminococcus, Lactobacillus and Prevotella. Nevertheless, it selectively encriched the Bacteroides, CF231, Echerichia, Alitispes, Butyrimous and Enterococcus (Figure 6).

**Figure 6.**
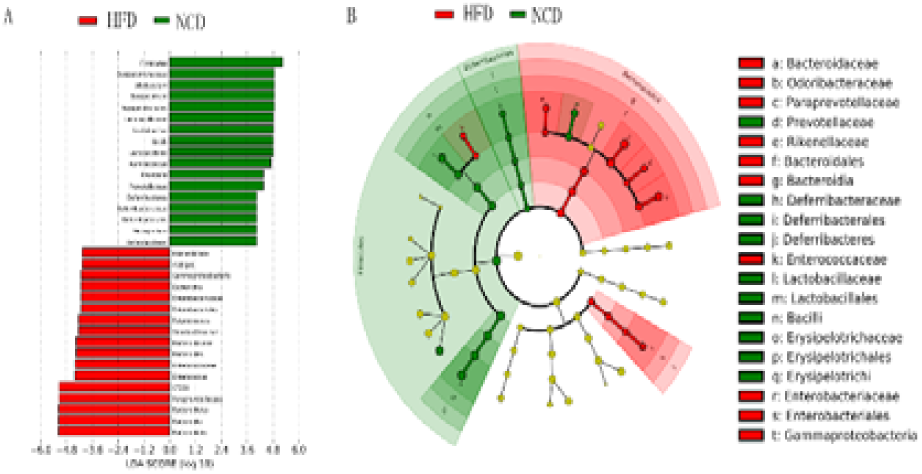
The significantly different species using the nonparametric factorial Kruskal-Wallis rank sum test at a significance level of 0.05. A: LAD score, an LDA score higher than 2 indicated a higher relative abundance in the corresponding group than in the other group; B: LEfSe taxonomic cladogram; different colors suggested enrichment of certain taxa in HFD and NCD groups.

Furthermore, the FHD vs HFD+ATO, and FHD vs HFD+JQII-5, HFD+JQI-7, HFD+LLY-606 or HFD+PC-26 groups were compared to analyze effects of the co-administration of ATO or probiotic strains on the HFD disturbed gut microbiota. The most obviously common reversion by ATO and all probiotic groups, was on the uncharacterized genus CF231 (**Figure 7A**). This genus was induced by HFD, from less than 0.0001% in NCD to 10.3% in HFD, and it was suppressed by co-administration of ATO or any of the tested probiotic strains. Another common reversed genus in all groups was *Bacteriodes* (**Figure 7B**). These common reversion on HFD induced dysbiosis of the gut microbiota indicated common effects of ATO and tested probiotics strains.

However, there are specificity in both HFD+ATO and HFD+probiotics groups. The most obvious difference between ATO and probiotic strains is determined by *Rumocuccous* and *Allobaculum*. *Rumococcus* was reversed by ATO but not by any of the probiotics (**Figure 7D**), but *Allobaculum* (**Figure 7C**) was reversed by all probiotic strains but not by ATO. Furthermore, *Allobaculum* was also one of the most dramatically changed genus, which is 7.57% in HFD, 20.79% in JQI-7, 13.85% in JQII-5, 18.29% in LLY-606 and 22.07% in PC-26.

**Figure 7.**
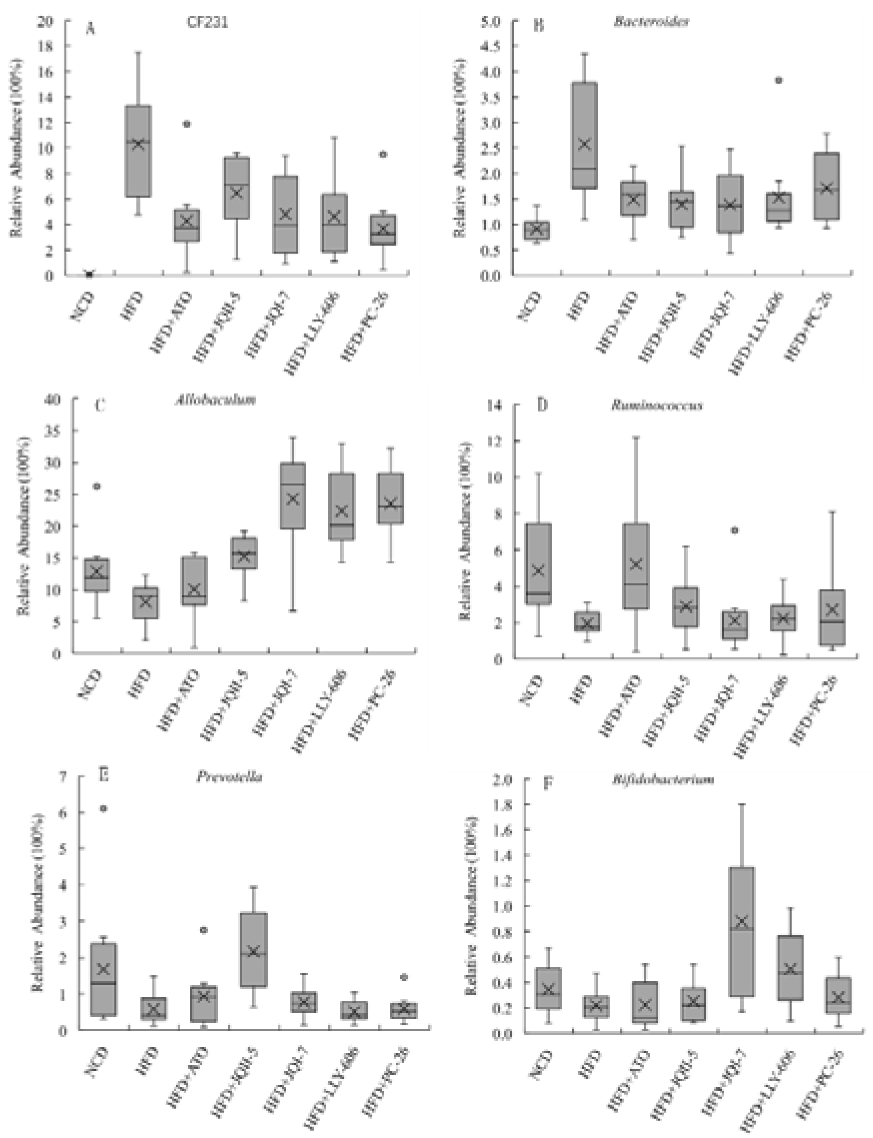
Relative abundance of key phylotypes in the gut modulated by ATO or probiotic strains.

In addition to commonly reversed genus, the specific altered genus by individual probiotic strains was analyzed. As shown in Figure 7E, the HFD suppressed *Preveotella* was reversed by JQII-5, but not by any other probiotic strains. And *Bifidobacterium* was enriched only by co-administration of JQI-7 (**Figure 7F**).

### 3.7 Effect of Different Probiotics on Lactic Acid and SCFAs in feces

We determined the fecal concentration of SCFAs, including acetic acid, propionic acid, butyric acid and lactate. The results indicated that HFD significantly decreased the fecal concentration of all tested acids. ATO co-administration reversed the decreased acetate and propionate, but not lactate and butyrate (**Figure 8**). All probiotic strains elevated acetic acid propionate and lactate, but the elevated butyrate was only seen in JIQ-7 and JQII-5.

**Figure 8.**
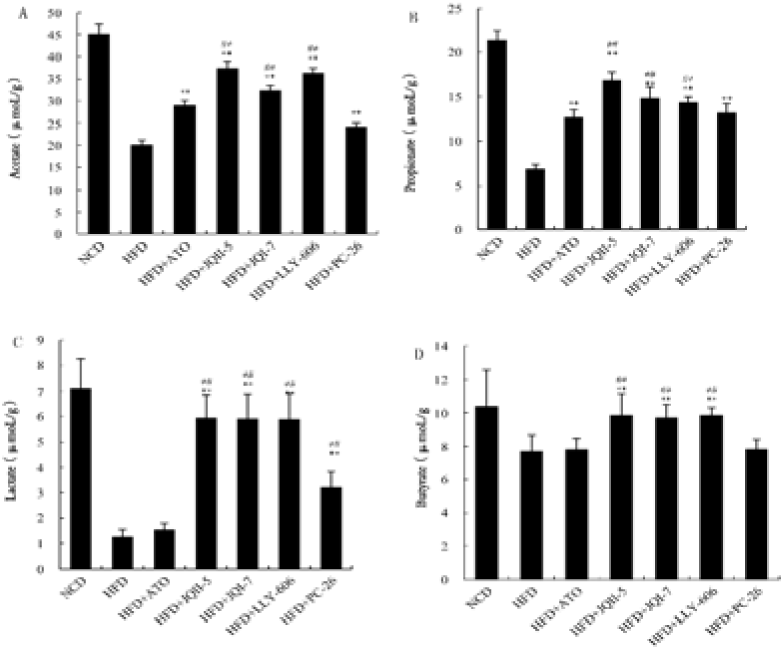
Concentration of fatty acid in fecal of golden hamsters. A: Acetate; B: Propionate; C: Lactate; D: Butyrate; ** means significant from

### 3.8 Correlation of key phylotypes and blood indicates

To demonstrated whether the key communities of gut microbiota were correlated to hypercholesterolemia, we performed an association analysis by pooling all groups together. In this part we mainly focused on phylotypes that were alerted by HFD and especially those that were reversed by ATO or probiotic strain.

As shown in **Figure 9**, genus enriched by HFD, such as CF231, *Bacteroides, Alitipes* and *Butricimounas* showed positive correlation with IL-1β and INS. The CF231 also showed significant positive correlation to TC (*p*<0.01) and TG (*p*<0.01).

**Figure 9.**
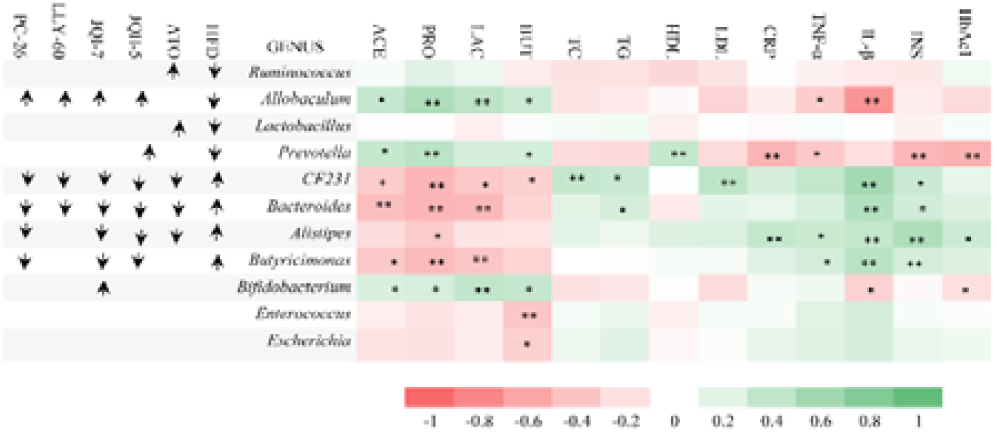
Correlations of key phylotypes to tested indexes related to cholesterolemia in golden hamsters. The color red indicates a negative correlation, whereas green shows a positive correlation. ↑ means: the phylotype in the corresponding row is increased in the group in the corresponding column.↓ means: the phylotype in the corresponding row is decreased in the group in the corresponding column.*means the significant level of p<0.05,**means the significant level of p<0.01.

*Allobaculum*, which was reversed by all probiotic strains but not by ATO, showed negative correlation to TNF-α (*p*<0.05) and IL-1β (*p*<0.01). However, what is unexpected is that the ATO altered *Rumococcus* showed no significant correlations to any tested factors. Another interesting observation was that, JQII-5 uniquely reversed genus *Prevotella* showed significant negative correlation to CRP (p<0.01), INS (*p*<0.01) and HbAc1 (*p*<0.01), showing a reflection of the effectiveness of JQII-5 on hypercholesterolemia observed above (**Figure 2**).

### 3.9 Correlation among key phylotypes

Then the correlations among the key phylotypes were analyzed. *Allococcum*, which was reversed by all probiotic strains but not by ATO, showed significantly negative correlation to almost all HFD induced genus, including CF231 (*p*<0.01), *Bacteroides* (*p*<0.01), and *Butricimonas* (*p*<0.01), *Enterococcus* (*p*<0.05) and *Escherichia* (*p*<0.05) (**Figure 10**). And it also seemed that all the HFD induced genus showed positive correlation to each other. These results indicated an indubitable role of *Allobacum* in the improvement of disturbed gut microbiota by HFD. Besides, *Allobacum* was also the sole correlated genus to *Bifidobacterium* (*p*<0.05). However, *Rumococcus*, which was reversed by ATO but not by any of the probiotic strains, was not correlated to any HFD altered genus.

**Figure 10.**
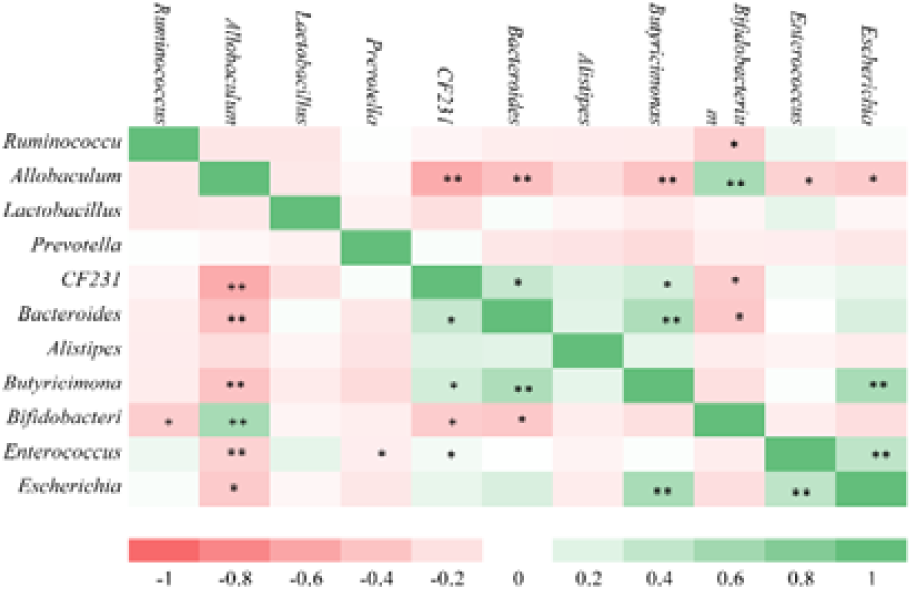
Correlation relationships among of key phylotypes.*means the significant level of p<0.05,**means the significant level of p<0.01.

## 4. Discussion

In this study, we found that compared with ATO, all tested probiotic strains showed mild effectiveness on both body weight and hypercholesterolemia of high-fat diet feed golden hamsters. However, after 8 weeks of administration, JQII-5, JIQ-7 and LLY-606 restored the HFD induced TC and LDL to a level comparable with ATO. ATO is a fully proofed and commonly prescribed cholesterol-lowering drug. The comparable effects of these 3 probiotic strains with ATO indicated full potential as a dietary supplementation for hypercholesterolemia. In fact, the effects of probiotic strains on hypercholesterolemia has been observed in many studies.^4, 6, 20, 21^ Yet, there is no doubt that vast of further researches are needed and there is still a long way to go.

To provide more solid evidence for their effectiveness on hypercholesterolemia, some factors related to dyslipidemia that are risky to health were also evaluated in this study. Inflammatory is a related state of hypercholesterolemia. TNF-α and IL-1β are well-known markers of inflammation and they have been reported to be elevated in hypercholesterolemic state. Some probiotic strains have been reported to reduce the expression of inflammation cytokines. ^22-25^ In the present study, the effectiveness of the tested strains on TNF-α and IL-1β were significant but not as effective as ATO. However, a comparable effect with ATO on CPR was achieved by co-administration of JQII-5 or JQI-7. CRP is a well-known marker of inflammation and it was a dyslipidemia induced risk factor for atherosclerosis, being used as a clinical parameter. ^26^ The effectiveness on CRP means further potential on improving syndrome induced by hypercholesterolemia.

Hypercholesterolemia is often accompanied by hyperglycemia, which is also proved in this study by elevated INS and HbA1c in HFD group (**Figure 4**). ^22^ And surprisingly, different from the mild effect on dyslipidemia, JQII-5 showed excellent effects on both INS and HbA1c as well as ATO. Higher levels of HbA1c was induced by long term and continuous dysglycemia, and higher levels of INS was an indicator of metabolic disorder in both lipid and glucose.^27^ The improvement of JQII-5 means an early influence on regulation of metabolic disorder induced by high-fat diet. From the comprehensive comparation of ATO with tested probiotic strains, significant, but retarded and mild effectiveness on both body weight, hypercholesterolemia, and inflammation was achieved by probiotic strain. And distinguishing effects on various index was found among the 4 probiotic strains. To provide some possible explanation for their shared and unique effects, the gut microbiota was analyzed. Just as expected, the microbiota structure after co-administration of ATO is markedly different from that of all probiotics strains groups, and the probiotic strain JQII-5 is different from the other 3 strains, indicating the different effects of ATO from probiotic strains and different effects of JQII-5 from the other 3 probiotic strains.

As more and more studies demonstrated the important influence of key communities in the gut microbiota,^28^ we investigated the HFD disturbed and ATO or individual probiotic strain reversed key phylotypes at the level of genus. The most obviously common reversion by ATO and all probiotic groups, was CF231 and *Bacteriodes*. CF231 was an uncharacterized genus, but it has been reported to be induced by HFD and suppressed by ATO in a previous study just as shown in the present study.^15^ And positive correlation of CF231 with TC, TC, LDL, IL-1β,and INS was found in this study, indicating its deleterious role in gut microbiota. *Bacteroides* was found to be more prevalent in western populations who consume higher animal-based diets compared with non-industrialized populations whose diets contain more dietary fibers.^29, 30^ A study including 165 older individuals found that higher levels of *Bacteroides* co-abundance group coincided with increased levels of CRP.^31^ These shared reversion on HFD induced CF231 and *Bacteroides* indicated an essential and effective role of ATO and tested probiotics strains.

*Ruminococcus* is a genus reversed uniquely by ATO and its reversion by ATO was also reported in a previous study. ^15^ Lower levels of *Ruminococcus* are associated with higher levels of CRP and IL-6. However, another study found that the abundance of *Rumicoccus gnavus* was higher in ACDV patients compared with controls. ^32^ In the present study, *Rumicoccus* was not found to be significantly correlated to any texted index (**Figure 9**), nor to almost any other genus (**Figure 10**). Whether the abundance changing of *Ruminococcus* is just a passive effect by ATO, and whether the ATO reversion of *Ruminococcus* would favor gut microbiota to benefit the hypercholesterolemia was still dubious.

One of the most amazing finding in this study is *Allobacum*, which is reversed by all tested probiotic strains and this effect was not shared by ATO. In a previous study, the relative abundance of *Allobacum* was increased by administration of bitter melon powder or by berberine, and the alteration of gut microbiota resulted in improvement of metabolic status in high fat diet-induced obese rats. ^33, 34^ In fact, high relative abundance of *Allobacum* was reported to be correlated to high H_2_ concentration in the gut and the portal.^35^ And H_2_ derived from fermentation of saccharides by gut microbiota was reported to suppress heptaic oxidative stress and adipose inflammatory in rats. ^36, 37^ As oxidative stress was assumed to be an important mechanism initiating inflammatory in obesity and hypercholesterolemia, the probiotic reversing *Alloculum* may improve the oxidative state of body and/or liver, which then help improve the inflammatory state of the body demonstrated in this study. Corresponding to this, a probiotic strain was reported to protect liver injury in rats via anti-oxidative and anti-inflammatory capacity. ^25^ Besides, different from the ATO induced *Ruminococcus, Alloculum* seemed to have significant negative correlation to most of the HFD induced genus, and positive correlation to *Bifidobacterium*. This genus commonly promoted by probiotics showed full chance to favor its host, and its favor is very probably done via favoring the gut microbiota.

In addition to the commonly promoted genus by all probiotic strains, JQII-5 also promoted *Prevotalla*. *Prevotalla* is associated with plant-rich diets, ^29^ and its suppression in this study was also supposed to be resulted from the high-fat and relatively low carbohydrates in HFD. Individuals with obesity have a lower abundance of *Prevotella* species in their gut. ^38^ Increased abundance of certain *Prevotella* species was reported to be associated with low-grade inflammation in systemic diseases, such as rheumatoid arthritis.^39^ *Prevotalla copri* was found to improve glucose metabolism and insulin sensitivity by a mechanism associated with fermentation of carbohydrates. ^40^ Consistent with these previous studies, co-administration of JQII-5 was found to be correlated with both inflammatory and glucose metabolism in the present studies (Figure 3 and Figure 4).

## 5. Conclusions

A comprehensive comparation of the ATO and 4 probiotic strains on high-fat induced hypercholesterolemia was done this study. Compared with ATO, probiotic strains were less effective on body weight, hypercholesterolemia, and inflammation. However, probiotics exert additional favorable effects on gut microbiota, making them excellent complements to cholesterol-lowering drugs like ATO. And these favorable effects were probably made by promoting key phenotypes, such as *Allobaculum* in gut microbiota, which may alleviate oxidative stress. However, the interaction between probiotic strains and targeted genus in gut microbiota is waiting to be confirmed in future study.

## Author contributions

Conceive and designed the experiments: Sudun, Gengyun Zhang, Shancen Zhao

Performed the experiments: Xiaofen He, Can Peng

Analyzed the data: Lifeng Liang, Songling Liu, Chen Xiao

Wrote the manuscript: Songling Liu, Chen Xiao, Sudun

## Acknowledgements

This study was supported by Science, Technology and Innovation Commission of Shenzhen Municipality (No.JCYJ20170817151208611) and Shenzhen Dapeng District Government (No. KY20180108).

## References

1 WHO Study Group, World Health Organ, Diet, nutrition, and the prevention of chronic diseases, Rep. Ser., 1990, 797, 1–204.

2 S. Yusuf, S. Hawken and S. Ounpuu, Effect of potentially modifiable risk factors associated with myocardial infarction in 52 countries (the INTERHEART study): case-control study (vol 364, pg 937, 2004), Lancet, 2004, 364, 1938–1938.

3 L. G. Ooi and M. T. Liong, Cholesterol-Lowering Effects of Probiotics and Prebiotics: A Review of in Vivo and in Vitro Findings, Int. J. Mol. Sci., 2010, 11, 2499–2522.

4 Z. Guo, X. M. Liu, Q. X. Zhang, Z. Shen, F. W. Tian, H. Zhang, Z. H. Sun, H. P. Zhang and W. Chen, Influence of consumption of probiotics on the plasma lipid profile: A meta-analysis of randomised controlled trials, Nutrition Metabolism and Cardiovascular Diseases, 2011, 21, 844–850.

5 Y. Bao, Z. Wang, Y. Zhang, J. Zhang, L. Wang, X. Dong, F. Su, G. Yao, S. Wang and H. Zhang, Effect of Lactobacillus plantarum P-8 on lipid metabolism in hyperlipidemic rat model, European Journal of Lipid Science and Technology, 2012, 114, 1230–1236.

6 T. P. Singh, R. K. Malik, S. G. Katkamwar and G. Kaur, Hypocholesterolemic effects of Lactobacillus reuteri LR6 in rats fed on high-cholesterol diet, Int. J. Food Sci. Nutr., 2015, 66, 71–75.

7 K. A. Greany, J. A. Nettleton, K. E. Wangen, W. Thomas and M. S. Kurzer, Probiotic consumption does not enhance the cholesterol-lowering effect of soy in postmenopausal women, J. Nutr., 2004, 134, 3277–3283.

8 K. L. Lindsay, M. Kennelly, M. Culliton, T. Smith, O. C. Maguire, F. Shanahan, L. Brennan and F. M. McAuliffe, Probiotics in obese pregnancy do not reduce maternal fasting glucose: a double-blind, placebo-controlled, randomized trial (Probiotics in Pregnancy Study), Am. J. Clin. Nutr., 2014, 99, 1432–1439.

9 M. T. Liong, Probiotics: A critical review of their potential role as antibypertensives, immune modulators, hypocholesterolemics, and perimenopausal treatments, Nutr. Rev., 2007, 65, 316–328.

10 J. R. Araujo, J. Tomas, C. Brenner and P. J. Sansonetti, Impact of high-fat diet on the intestinal microbiota and small intestinal physiology before and after the onset of obesity, Biochimie, 2017, 141, 97–106.

11 I. M. Indias, F. Cardona, F. J. Tinahones and M. I. Queipo-Ortuno, Impact of the gut microbiota on the development of obesity and type 2 diabetes mellitus, Front. Microbiol., 2014, 5, 1–10.

12 M. G. Rooks, P. Veiga, L. H. Wardwell-Scott, T. Tickle, N. Segata, M. Michaud, C. A. Gallini, C. Beal, J. E. T. van Hylckama-Vlieg, S. A. Ballal, X. C. Morgan, J. N. Glickman, D. Gevers, C. Huttenhower and W. S. Garrett, Gut microbiome composition and function in experimental colitis during active disease and treatment-induced remission, Isme Journal, 2014, 8, 1403–1417.

13 A. N. Ananthakrishnan, C. Luo, V. Yajnik, H. Khalili, J. J. Garber, B. W. Stevens, T. Cleland and R. J. Xavier, Gut Microbiome Function Predicts Response to Anti-integrin Biologic Therapy in Inflammatory Bowel Diseases, Cell Host & Microbe, 2017, 21, 603–610.

14 J. H. Yang, P. Bhargava, D. McCloskey, N. Mao, B. O. Palsson and J. J. Collins, Antibiotic-Induced Changes to the Host Metabolic Environment Inhibit Drug Efficacy and Alter Immune Function, Cell Host & Microbe, 2017, 22, 757–765.

15 T. J. Khan, Y. M. Ahmed, M. A. Zamzami, S. A. Mohamed, I. Khan, O. A. S. Baothman, M. G. Mehanna and M. Yasir, Effect of atorvastatin on the gut microbiota of high fat diet-induced hypercholesterolemic rats, Sci. Rep., 2018, 8.662

16 Y. Yu, Q. Liu, H. Li, C. Wen and Z. He, Alterations of the Gut Microbiome Associated With the Treatment o f Hyperuricaemia in Male Rats, Front. Microbiol., 2018, 9, 2233.

17 S. Das, A. Datta, C. Bagchi, S. Chakraborty, A. Mitra and S. K. Tripathi, A Comparative Study of Lipid-Lowering Effects of Guggul and Atorvastatin Monotherapy in Comparison to Their Combination in High Cholesterol Diet-Induced Hyperlipidemia in Rabbits, J. Diet. Suppl., 2016, 13, 495–504.

18 C. Peng, Y. Li, Y. Chen, Y. Kong, Y. Lin, H. Li, L. Chen, Screening and comparison of cholesterol-lowing Lactobacillus strains, Chinese J. Microecology, 2017, 29, 249–255.

19 C. Wang,P. X. Gao, J. N. Xu, W. G. Yu, X. Z. Lu, Establishment and Application of Extraction and Determination Method for Short-Chain Fatty Acids in Feces of Mice, Progress in Modern Biomedicine, 2017, 17, 1012–1015.

20 X. Zhao, F. Higashikawa, M. Noda, Y. Kawamura, Y. Matoba, T. Kumagai and M. Sugiyama, The Obesity and Fatty Liver Are Reduced by Plant-Derived Pediococcus pentosaceus LP28 in High Fat Diet-Induced Obese Mice, PLoS One, 2012, 7, e30696.

21 F. T. u. Lim, S. M. Lim and K. Ramasamy, Cholesterol lowering by Pediococcus acidilactici LAB4 and Lactobacillus plantarum LAB12 in adult zebrafish is associated with improved memory and involves an interplay between npc1l1 and abca1, Food Funct., 2017, 8, 2817–2828.

22 X. Zhou, W. Zhang, X. Liu, W. Zhang and Y. Li, Interrelationship between diabetes and periodontitis: Role of hyperlipidemia, Arch. Oral Biol., 2015, 60, 667–674.

23 N. Esser, N. Paquot and A. J. Scheen, Anti-inflammatory agents to treat or prevent type 2 diabetes, metabolic syndrome and cardiovascular disease, Expert Opinion on Investigational Drugs, 2015, 24, 283–307.

24 W. Jiang, Y. Cen, Y. Song, P. Li, R. Qin, C. Liu, Y. Zhao, J. Zheng and H. Zhou, Artesunate attenuated progression of atherosclerosis lesion formation alone or combined with rosuvastatin through inhibition of pro-inflammatory cytokines and pro-inflammatory chemokines, Phytomedicine, 2016, 23, 1259–1266.

25 Y. Wang, Y. Li, J. Xie, Y. Zhang, J. Wang, X. Sun and H. Zhang, Protective effects of probiotic Lactobacillus casei Zhang against endotoxin- and D-galactosamine-induced liver injury in rats via anti-oxidative and anti-inflammatory capacities, Int. Immunopharmacol., 2013, 15, 30–37.

26 D. Medenwald, S. Dietz, D. Tiller, A. Kluttig, K. Greiser, H. Loppnow, J. Thiery, S. Nuding, M. Russ, A. Fahrig, J. Haerting and K. Werdan, Inflammation and echocardiographic parameters of ventricular hypertrophy in a cohort with preserved cardiac function, Open heart, 2014, 1, e000004–e000004.

27 B. B. Kahn and J. S. Flier, Obesity and insulin resistance, J. Clin. Invest., 2000, 106, 473–481.

28 L. Zhao, F. Zhang, X. Ding, G. Wu, Y. Y. Lam, X. Wang, H. Fu, X. Xue, C. Lu, J. Ma, L. Yu, C. Xu, Z. Ren, Y. Xu, S. Xu, H. Shen, X. Zhu, Y. Shi, Q. Shen, W. Dong, R. Liu, Y. Ling, Y. Zeng, X. Wang, Q. Zhang, J. Wang, L. Wang, Y. Wu, B. Zeng, H. Wei, M. Zhang, Y. Peng and C. Zhang, Gut bacteria selectively promoted by dietary fibers alleviate type 2 diabetes, Science, 2018, 359, 1151–1156.

29 C. De Filippo, D. Cavalieri, M. Di Paola, M. Ramazzotti, J. B. Poullet, S. Massart, S. Collini, G. Pieraccini and P. Lionetti, Impact of diet in shaping gut microbiota revealed by a comparative study in children from Europe and rural Africa, Proc. Natl. Acad. Sci. U. S. A., 2010, 107, 14691–14696.

30 S. L. Schnorr, M. Candela, S. Rampelli, M. Centanni, C. Consolandi, G. Basaglia, S. Turroni, E. Biagi, C. Peano, M. Severgnini, J. Fiori, R. Gotti, G. De Bellis, D. Luiselli, P. Brigidi, A. Mabulla, F. Marlowe, A. G. Henry and A. N. Crittenden, Gut microbiome of the Hadza hunter-gatherers, Nature Communications, 2014, 5.

31 T. Yatsunenko, F. E. Rey, M. J. Manary, I. Trehan, M. G. Dominguez-Bello, M. Contreras, M. Magris, G. Hidalgo, R. N. Baldassano, A. P. Anokhin, A. C. Heath, B. Warner, J. Reeder, J. Kuczynski, J. G. Caporaso, C. A. Lozupone, C. Lauber, J. C. Clemente, D. Knights, R. Knight and J. I. Gordon, Human gut microbiome viewed across age and geography, Nature, 2012, 486, 222–227.

32 Z. Jie, H. Xia, S.-L. Zhong, Q. Feng, S. Li, S. Liang, H. Zhong, Z. Liu, Y. Gao, H. Zhao, D. Zhang, Z. Su, Z. Fang, Z. Lan, J. Li, L. Xiao, J. Li, R. Li, X. Li, F. Li, H. Ren, Y. Huang, Y. Peng, G. Li, B. Wen, B. Dong, J.-Y. Chen, Q.-S. Geng, Z.-W. Zhang, H. Yang, J. Wang, J. Wang, X. Zhang, L. Madsen, S. Brix, G. Ning, X. Xu, X. Liu, Y. Hou, H. Jia, K. He and K. Kristiansen, The gut microbiome in atherosclerotic cardiovascular disease, Nature Communications, 2017, 8, 845–856.

33 X. Zhang, Y. Zhao, M. Zhang, X. Pang, J. Xu, C. Kang, M. Li, C. Zhang, Z. Zhang, Y. Zhang, X. Li, G. Ning and L. Zhao, Structural Changes of Gut Microbiota during Berberine-Mediated Prevention of Obesity and Insulin Resistance in High-Fat Diet-Fed Rats, PLoS One, 2012, 7, e42529.

34 J. Bai, Y. Zhu and Y. Dong, Modulation of gut microbiota and gut-generated metabolites by bitter melon results in improvement in the metabolic status in high fat diet-induced obese rats, J. Funct. Foods, 2018, 41, 127–134.

35 N. Nishimura, H. Tanabe, E. Komori, Y. Sasaki, R. Inoue and T. Yamamoto, Transplantation of High Hydrogen-Producing Microbiota Leads to Generation of Large Amounts of Colonic Hydrogen in Recipient Rats Fed High Amylose Maize Starch, Nutrients, 2018, 10, 144.

36 N. Nishimura, H. Tanabe, Y. Sasaki, Y. Makita, M. Ohata, S. Yokoyama, M. Asano, T. Yamamoto and S. Kiriyama, Pectin and high-amylose maize starch increase caecal hydrogen production and relieve hepatic ischaemia-reperfusion injury in rats, Br. J. Nutr., 2012, 107, 485–492.

37 N. Nishimura, H. Tanabe, M. Adachi, T. Yamamoto and M. Fukushima, Colonic Hydrogen Generated from Fructan Diffuses into the Abdominal Cavity and Reduces Adipose mRNA Abundance of Cytokines in Rats, J. Nutr., 2013, 143, 1943–1949.

38 R. E. Ley, P. J. Turnbaugh, S. Klein and J. I. Gordon, Microbial ecology - Human gut microbes associated with obesity, Nature, 2006, 444, 1022–1023.

39 J. M. Larsen, The immune response to Prevotella bacteria in chronic inflammatory disease, Immunology, 2017, 151, 363–374.

40 P. D. Cani, Human gut microbiome: hopes, threats and promises, Gut, 2018, 67, 1716–1725.

