## Supplementary figures and images for "Probiotic strains improve high-fat diet-induced hypercholesterolemia through modulating gut microbiota in ways different from atorvastatin"

### supplementary S1-S5

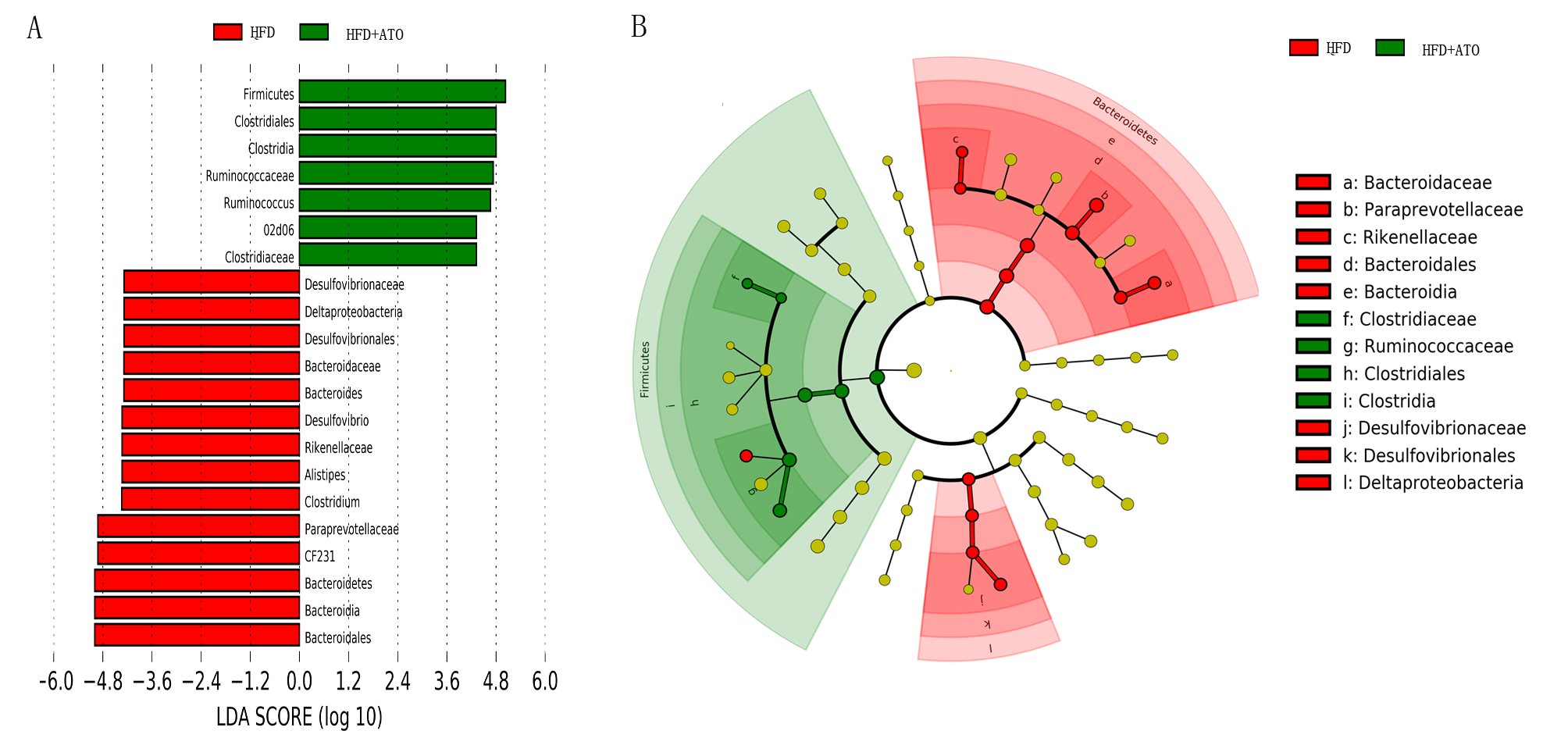

### supplementary S1-S5

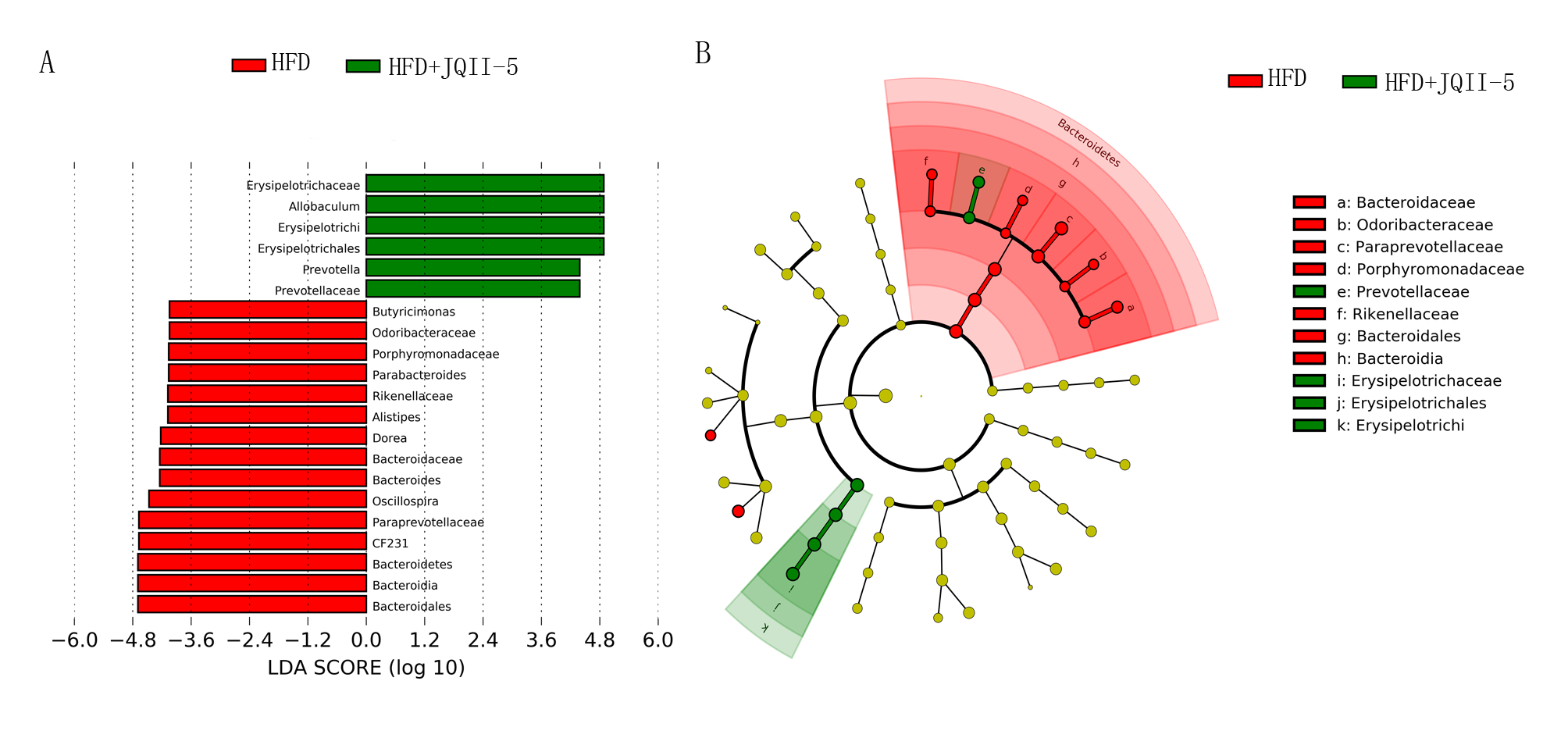

### supplementary S1-S5

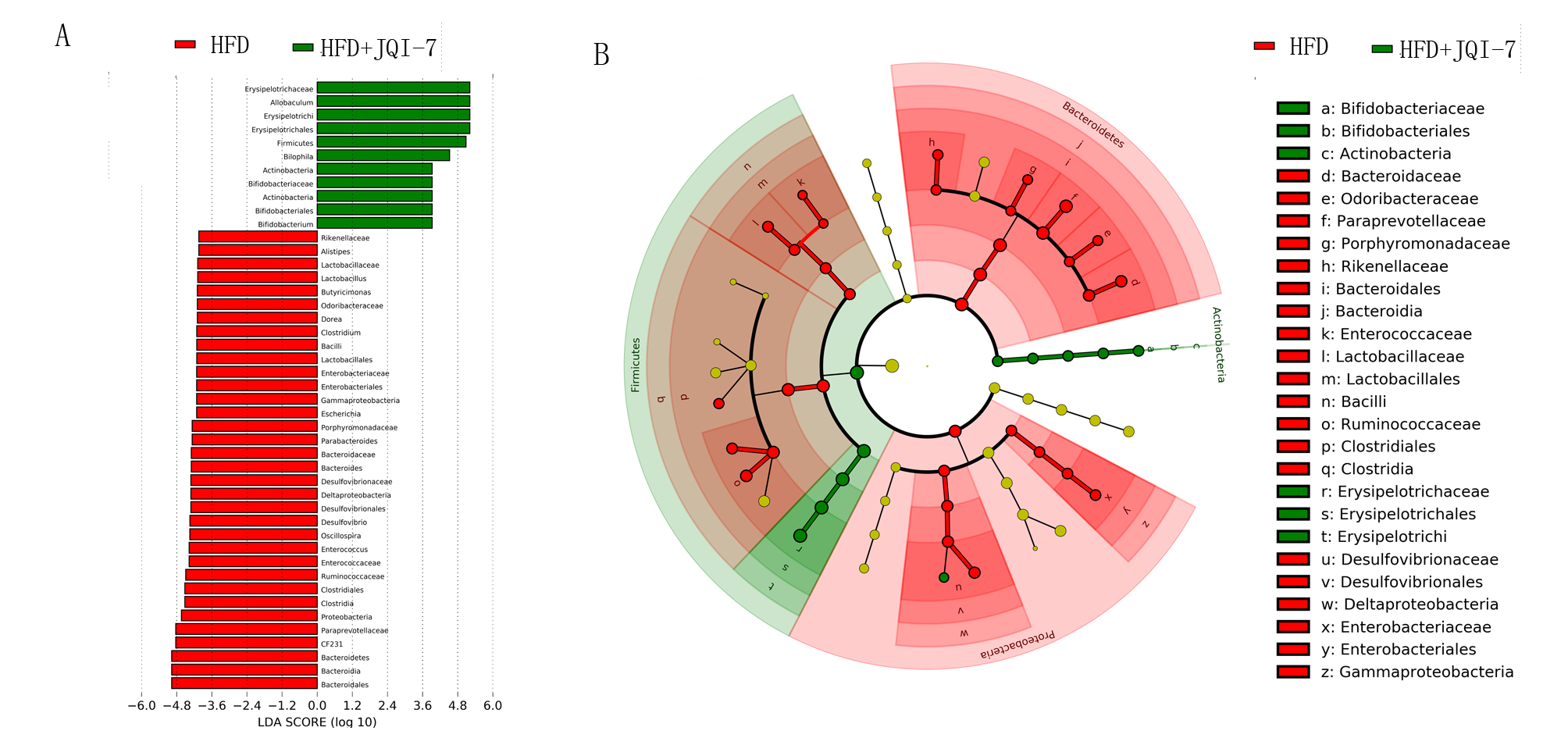

### supplementary S1-S5

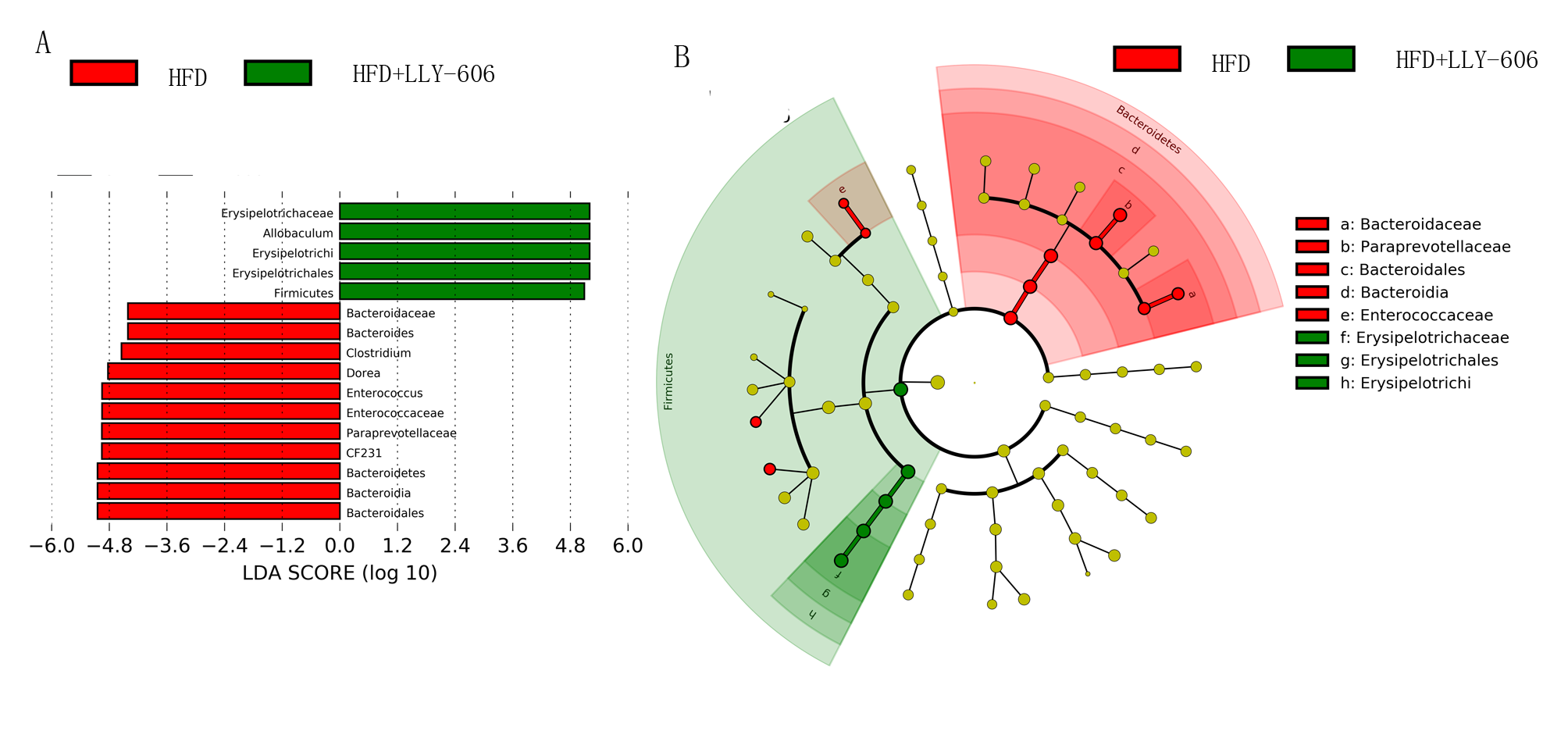

### supplementary S1-S5

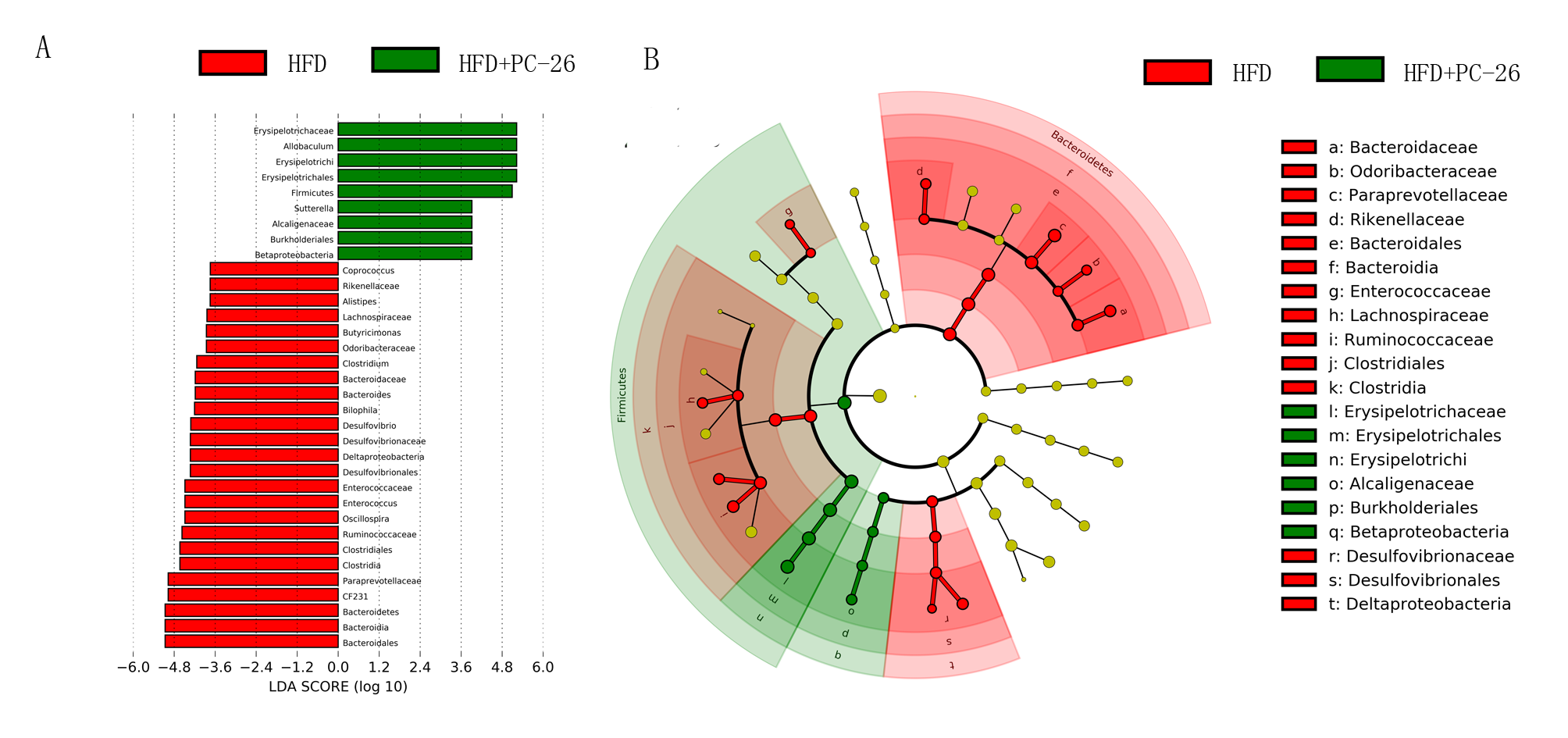
